# Spatiotemporal Mapping of Cherry Blossom Blooming by Semi-Automatic Observation System with Street-Level Photos

**DOI:** 10.1101/2023.04.13.536831

**Authors:** Narumasa Tsutsumida, Shuya Funada

## Abstract

The importance of floral phenology as a critical indicator of regional climate change and ecosystem services is widely recognized. The annual blooming of cherry blossoms is a nationally celebrated event in Japan, and historical phenological records have been used to document regional climate change. The cultural ecosystem services provided by this phenomenon are important as they not only signal the arrival of spring but also offer a picturesque spring landscape. Despite its importance, constructing a spatiotemporal record of cherry blossom blooming is challenging due to the limited coverage of traditional stationary observations. To address this issue, citizen-based observation programs and remote sensing applications have been implemented; nevertheless, these strategies are still limited by infrequent and insufficient observations throughout space and time. To compensate, we developed a flower detection model for geographically and temporally dispersed street-level photos that may be used as the core component of a semi-automatic observation system. Specifically, we developed a customized YOLOv4 model for cherry blossom detection from street-level photos obtained through Mapillary, one of the social sensing data repositories. The detection model achieved an overall accuracy, recall, and precision of 86.7%, 70.3%, and 90.1%, respectively. By using observation coordinates and dates attached to Mapillary photos, we mapped the probability of cherry trees blooming in a spatial grid of dimensions 10 m x 10 m on a daily basis. With sufficient observations, start, peak, and end of blooming were estimated through time series analysis. A case study conducted at Saitama University’s main campus in 2022 confirmed the possibility of mapping the presence of cherry blossoms and their blooming timing automatically. Since our approach relies solely on geotagged street-level photos that can be taken by anyone with no prior knowledge of cherry tree species identification, we anticipate that it will be easier to build blooming records over space and time than conventional stationary observations or citizen-based observation programs. This novel approach also has potential applications for detecting other species as well.

## 1. Introduction

Floral phenology (the timing and intensity of flowering), has recently gained recognition within the scientific community as a valuable indicator of both climate change and ecosystem services (Chen et al., 2019; Elzinga et al., 2007). However, accurately estimating flowering events can be a formidable challenge, given their regulation not only by local and regional environmental variables such as temperature, rainfall (El Yaacoubi et al., 2014; Grab and Craparo, 2011), and photoperiod (Bicknell and Jaksons, 2018), but also by factors such as water stress, soil condition (Ish-Shalom-Gordon, 1993), and genetic modulation (Elzinga et al., 2007). Because of the complex mechanisms underlying floral phenology, the establishment of a spatial and temporal record of flowering events is crucial to increase the understanding of its reaction patterns to shifts in local environmental factors (Chen et al., 2019).

Currently, extended *in-situ* observation projects are regarded as the most common method to record floral phenology on a global scale.

To monitor seasonal differences, the Japan Meteorological Agency (JMA) has annually conducted seasonal surveys of plants and animals since 1953 (Doi et al., 2021).

However, because of difficulties that have arisen in the observation of some species, together with inadequate funding, the project has suffered a severe decline since 2021 (Doi et al., 2021). For example, 40% of the targeted sites have been terminated, and 94% of phenological events, including both plants and animals, have ceased to be observed (Doi et al., 2021). Despite the importance of official long-term observations at particular sites, the spatiotemporal variability of floral phenology due to regional climate has never been taken into consideration.

To address the lack of observations across space and time, some supportive approaches have been considered as workable alternatives to regional phenological observations.

The efficacy of using data provided by non-professionals has been investigated in the field of citizen science. Taylor et al. (2019) examined this topic and demonstrated the usefulness of citizen science data in a case study. By using both statistical and process-based models, phenological events can be estimated based on both citizen science data and observational data. However, MacKenzie et al. (2017) noted the potential inaccuracy in plant species identification and citizen-reported locations. This highlights the complexity and need for caution when interpreting citizen-provided data.

Rosemartin et al. (2021) discussed a citizen science project and showed the importance of the sustainable nature of long-term citizen-based observations. These factors should be taken into account when considering the potential of citizen science data.

Satellite-based remote sensing (RS) techniques have also been widely utilized for floral phenological studies. Chen et al. (2019) employed an enhanced bloom index (EBI) in order to monitor the floral activity of almond (Prunus dulcis) orchards based on time-series multispectral RS images. Similarly, Dixon et al. (2021) proposed a machine-learning method to predict flowering events using time-series multispectral RS images. These RS studies have shown great potential for gathering and mapping floral phenological information over broad temporal and spatial scales. However, the challenges posed by model estimation error and the coarse temporal resolution of RS observations may result from infrequent revisits and cloud cover and should be addressed. Tackling such challenges is critical for an accurate inventory of floral phenology. Another approach is to create a robust object detection model for flower detection from a proximity sensing image. August et al. (2020) have implemented an innovative deep-learning model that accurately classifies various plant species and generates a species occurrence map from a dataset of over 23,000 Flicker photos.

Employing the inception architecture developed by a convolutional neural network (CNN), the model is capable of detecting 519 unique species in London, UK. Van Horn et al. (2017) and Garcin et al. (2021) have proposed an image database that can aid in the creation and validation of an efficient deep-learning model for species recognition and classification.

The Japanese cherry blossom season is a noteworthy event that has attracted attention from researchers who study floral phenology. Scholars acknowledge its importance not only in providing cultural ecological services (Katsuda et al., 2022; Nagai et al., 2019), but also in its long-term observational records, which can be used to better understand the impact of local climatic conditions. Long-term stationary observational data have been used by researchers to examine the relationship between the timing of cherry blossom blooming and the phenomenon of climate change. Several studies have been undertaken in this regard (Chung et al., 2009; Masago and Lian, 2022; Nagai, Morimoto, et al., 2020; Nagai, Saitoh, et al., 2020; Shi et al., 2017). Recent research has also used remote sensing to observe the temporal and spatial patterns of cherry blossom blooming (Hassan et al., 2015; Kim et al., 2022; Lim et al., 2020). However, no blooming records over time have been compiled using this approach. Some studies have focused on using social media, as blossoming events can be identified from this data without the need for careful interpretation (August et al., 2020; ElQadi et al., 2021; Horikawa et al., 2022; Morishita et al., 2015; Shin et al., 2022). Shin et al. (2022) employed Google Trends to analyze user interest in cherry blossoms through Google searches over time, demonstrating its usefulness in estimating cherry flowering phenology when observational data is lacking. ElQadi et al. (2021) used geotagged and text-tagged social sensing photos from Flickr, extracting cherry-related images from the attached labels. However, complex data pre-processing was required to eliminate mislabeled photos. A kernel density was introduced to reduce inaccurate location errors, and the estimated full bloom event was mapped at the city level. When developing an automatic cherry blossom detection system, Morishita et al. (2015) proposed using a car-mounted smartphone to record the blossoming trees. They developed a participatory sensing system that automatically extracts cherry blossoms blooming from videos recorded by the smartphone and then shares the information among users in quasi-realtime. To detect the blooming of cherry blossoms, Morishita et al. (2015) used histogram-based color analysis and region-based fractal dimension analysis to obtain multiple frames of the video. However, because a color histogram is species-specific, their approach, while useful for detecting flowering events, lacks scalability for detecting analogous events of other species. Another limitation is that this technique was not used to estimate cherry flowering phenology, which could have provided insight into how cherry flowering responds to local climate and environments.

Many studies support blossom detection and biodiversity monitoring using image analysis from multiple data sources, such as social media data and street-level imagery. These studies have highlighted the possible use of deep-learning models. None of these studies has yet been able to successfully record the spatiotemporal blooming of cherry blossoms at a local level.

This study endeavors to address the gaps in previous research by introducing an innovative approach for obtaining records of cherry blossom blooming phenology. The method utilizes geotagged street-level photographs, which are subsequently analyzed by a deep-learning model. This study focuses on the Yoshino cherry (Prunus × yedoensis, Someiyoshino) because it is an important indicator of climate change (Aono and Kazui, 2008; Aono and Saito, 2010) and cultural ecosystem services (Nagai et al., 2016). Our research demonstrates an innovative, semi-automatic method for compiling street-level photos that are geotagged to create chronicles of cherry blossom blooming.

## 2. Materials and Methods

### 2.1 Overview

The proposed methodological framework is illustrated in Figure 1. The comprehensive approach has three main constituents, namely the Data, Model, and Phenological analysis components. This streamlined workflow enables it to be used by a broad range of case studies and various species. The data component involves the use of street-level images captured by the authors and uploaded to Mapillary, which is one of the social sensing data platforms. This enhances the scope and applicability of our method by incorporating socially-sensed images and eliminates the need for on-site data collection specific to phenological observations. The Model component detects cherry blossom blooming and can be replaced by alternative object detection models if required. In this study, we used the YOLOv4 model (Bochkovskiy et al., 2020). This common object-detection model allows us to determine both the boundary and the probability of the target class. The detection probability is important in estimating floral phenological events, which are explained in detail in the following sections. Finally, the phenological analysis component maps the phenological records of the blooming phase. We aggregated the available photo-shooting points into a 10 m spatial grid and calculated the spatially-averaged daily probability to reduce potential geolocation error margins and distance gaps between observed points and cherry tree locations. Then, using a threshold approach, phenological events such as the start, peak, and end of blooming were analyzed from the time-series probability at each grid.

**Figure 1.**
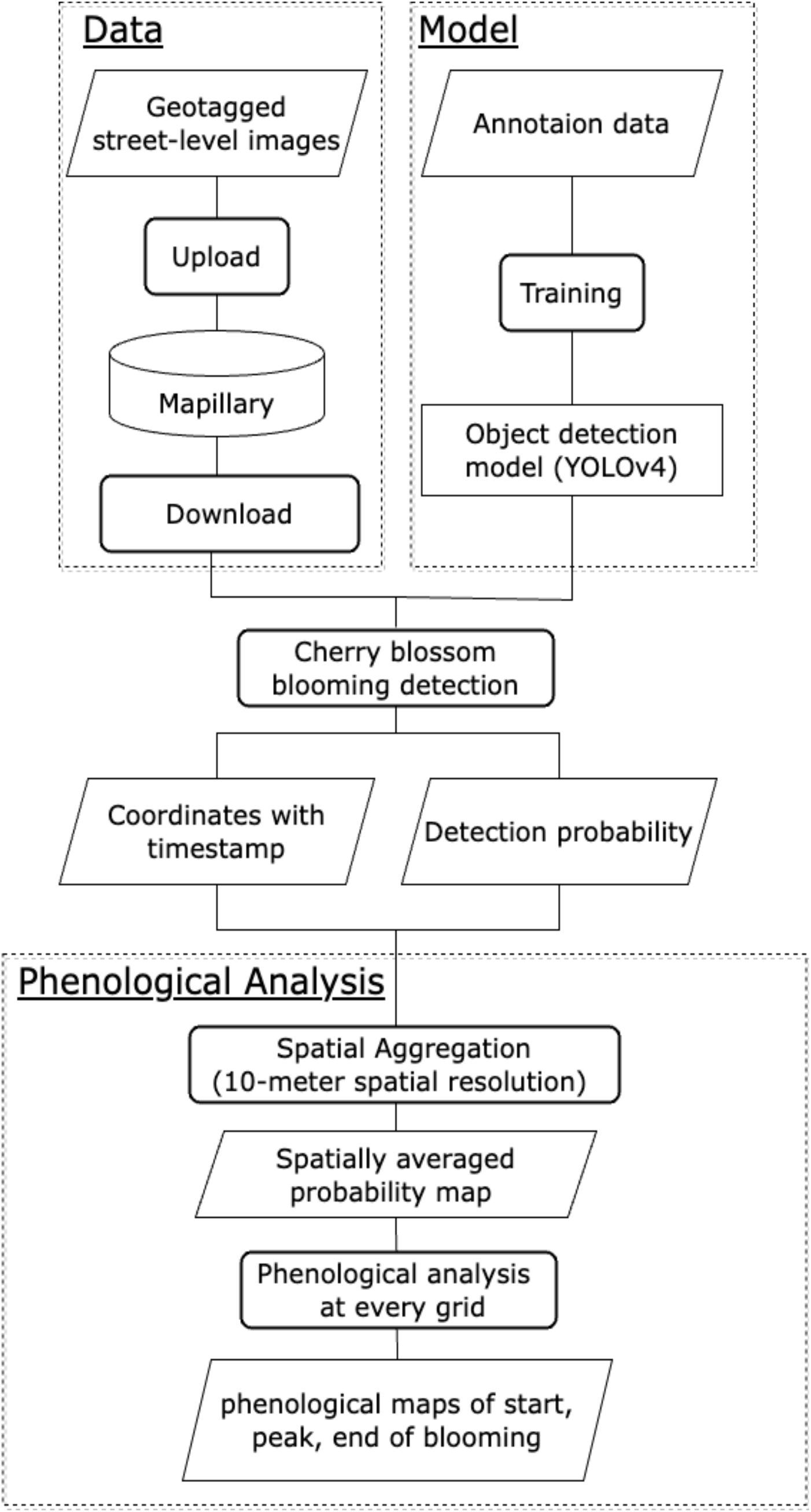
Flowchart of the study.

### 2.2 Data

The study area was the main campus of Saitama University, located in Japan. A GoPro 10 and iPhone 13 were used to capture the cherry blossoms blooming. A total of 7,168 geotagged street-level photos were collected (Figure 2). These photos were taken during the period between March 15 and April 10, 2022, as frequently as possible, with the exception of rainy days. The use of the time-lapse mode in the GoPro settings, as well as the Mapillary iOS app, allowed us to obtain a sequenced trajectory of the camera movement. The photos contained their respective coordinate information (specifically, latitude and longitude), photo angle, and observation date. These photos were uploaded to Mapillary, a platform designed to store geotagged street-level photos from all over the world. Since all photos on Mapillary are licensed under CC-BY-SA, users may download and use them as per the regulations. This platform was crucial to our research protocol since it improved the efficiency and scalability of our approach for gaining access to geotagged street-level photos for cherry blossom recognition.

**Figure 2.**
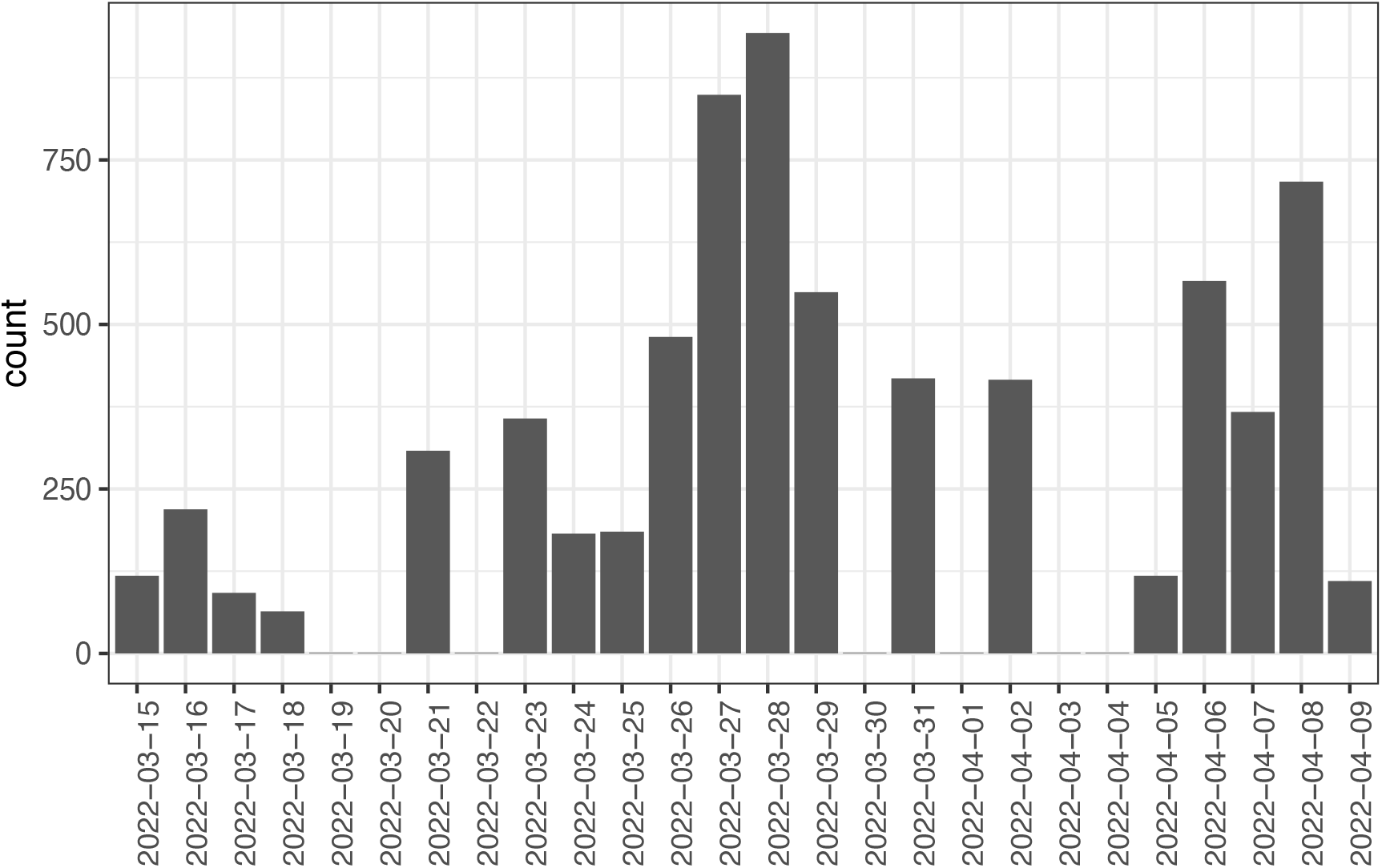
Number and dates of observations.

### 2.3 Model

The YOLOv4 model developed by Bochkovskiy et al. (2020) was used to identify cherry blossoms in street-level photographs. YOLOv4, a CNN-based object detection model, partitions the input images into multiple regions and delineates the bounding box surrounding the likelihood of the target class in each region. We selected YOLOv4 because it can operate with a single standard graphics processing unit to conduct training and prediction using large datasets. Further information regarding the model can be found in the article by Bochkovskiy et al. (2020). Wu et al. (2020) and Cheng and Zhang (2020) have previously introduced YOLOv4 for flower detection, with Wu et al. (2020) detecting apple flowers in smartphone photos captured at a distance of 30– 50 cm from trees and Cheng and Zhang (2020) detecting and categorizing 102 flower species from the Oxford 102 Flower dataset (Nilsback and Zisserman, 2008). However, their usage was restricted to singular flower images captured at close range.

Web-based repositories such as Flickr (https://www.flickr.com/) were sourced for 698 open-licensed photographs of cherry blossoms blooming that were then annotated and used to train the YOLOv4 model. Our focus was solely on the Prunus × yedoensis, commonly known as the Yoshino cherry tree, as it is highly favored and extensively planted throughout Japan.

### 2.4 Phenological analysis

In this section, we present a phenological analysis chain used to study the blooming of cherry trees. Initially, we combined available photo shooting points in a 10 m spatial grid. Although the geo-tagged street-level photos were taken with GPS-enabled GoPro and iPhone devices, the locational accuracy was not precise. To address this issue, we calculated the average detection probability for cherry blossoms in every grid with a 10 m × 10 m resolution. This approach helped to minimize coordination errors. Daily probability maps of cherry blossoms blooming during the study period were produced while considering temporal changes in the cherry blossom blooming probability at every grid. To further refine our analysis, we filled in the non-observational values of the averaged probability with a spline curve and applied a 7-day moving average to every grid. Lastly, we calculated the timing of cherry blossom blooming, defining the start and end of blooming as the first and last days when the averaged probability exceeded 0.25. The peak of blooming was defined as the day with the maximum averaged probability. These thorough analyses produced reliable information about cherry blossoming during the study period and highlighted the efficacy of the proposed methods.

## 3. Results

### 3.1 Building spatiotemporal flowering event records

The daily probability maps were useful for determining spatiotemporal flowering event records in the context of the spatiotemporal dynamics of cherry blossom blooming (Figure 3). The flowering period for Prunus × yedoensis began around 21 March 2022, with peak blooms observed around 31 March 2022. The disappearance of the blooms was recorded around 9 April 2022. Despite our intended focus on the blossoms of Prunus × yedoensis, we observed slight variations in the blooming timing and intensity across different spatial locations, which we attribute to localized environmental factors such as soil composition and photoperiod. Furthermore, the majority of observations were limited to prominent footpaths and major roads due to accessibility issues, and the frequency of observations was unevenly distributed across the established observation points. These inconsistencies and infrequent observations are a common challenge encountered during ecological field surveys.

**Figure 3.**
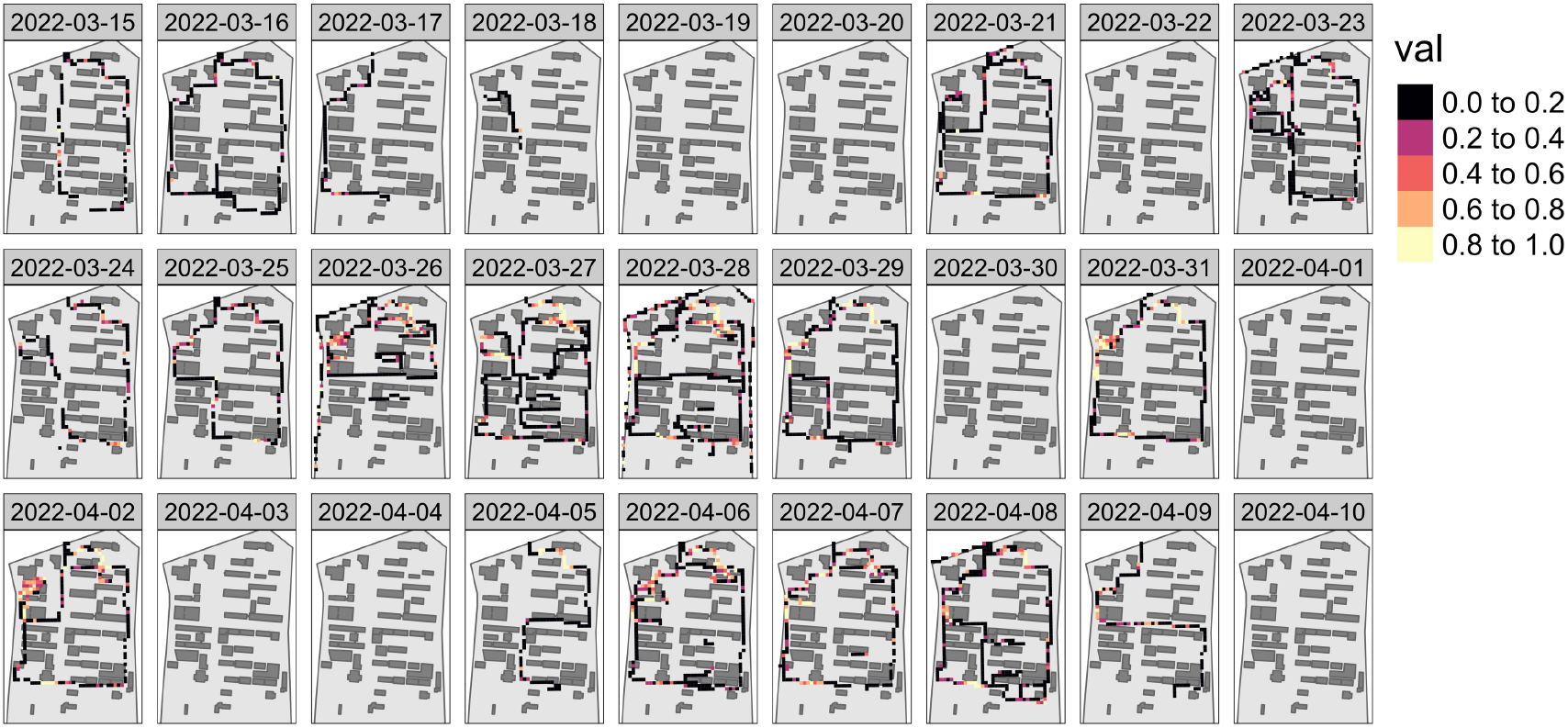
Daily maps of averaged cherry blossoms detection probability of cherry blooming in 10 m × 10 m spatial resolution.

The phenological records of cherry blossoms, including their start, peak, and end of blooming, are presented in Figure 4. The start, peak, and end of blooming were estimated on a limited number of grids, as indicated by the daily probability maps in Figure 3. This estimation was only possible in areas where an adequate number of photos of the cherry blossoms were taken. To update the blooming record, an automated detection method was used to identify five areas where Yoshino cherry trees were planted. Figure 4 depicts the local variation in the start, peak, and end of blooming in the study area. There were variations in the blooming times of the trees; the start, peak, and end of blooming were primarily observed between March 24th and 26th, March 29th and April 3rd, and April 5th and 6th, respectively. However, it is important to note that the start, peak, and end of blooming were not identical across all areas.

**Figure 4.**
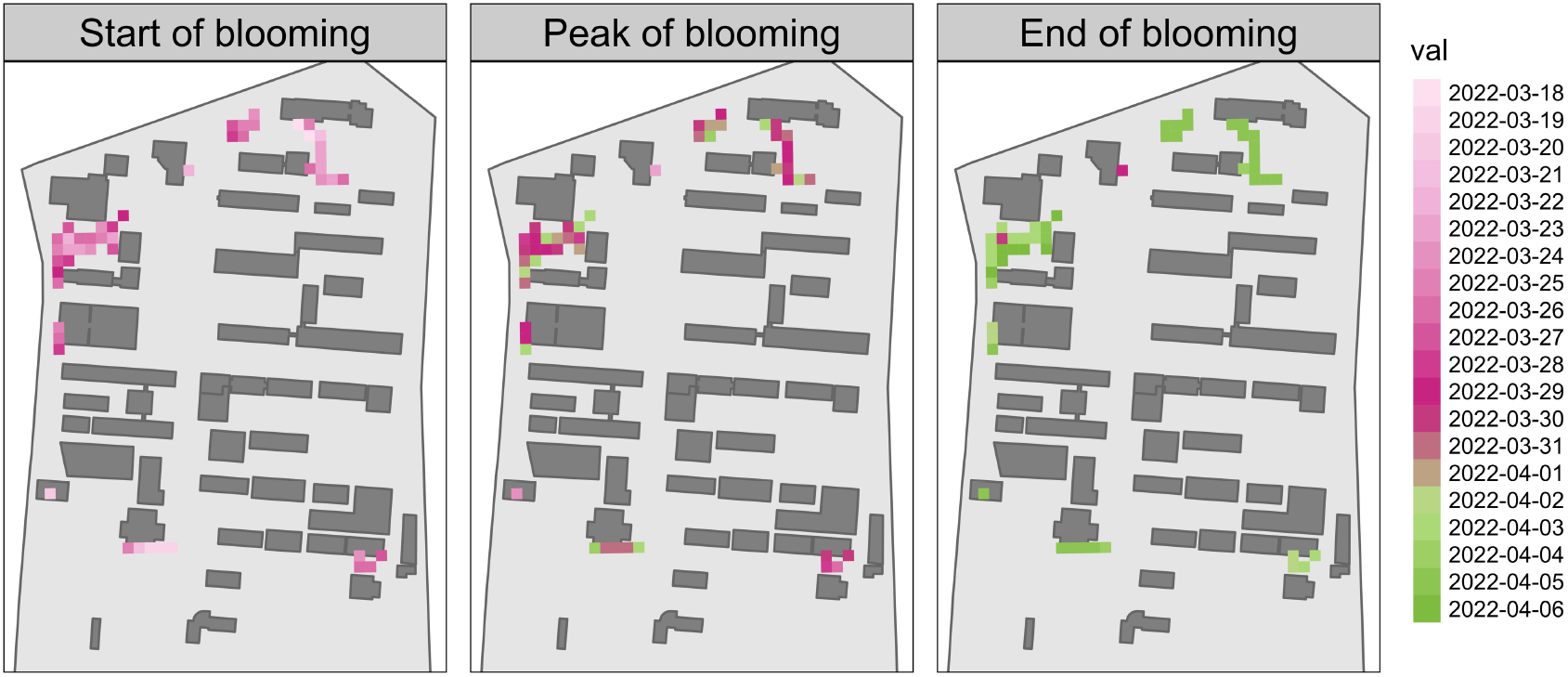
Dates of start, peak, and end of cherry blossom in 10 m × 10 m spatial resolution.

We investigated the relationship between the temporal change in cherry blossom blooming and its detection probability within a 10 m × 10 m spatial grid to thoroughly understand its flowering patterns during the study period (Figure 5). A total of 66 photos were found in the grid. The detection probability displayed a spatially averaged association with each blooming stage. The first time when the moving averaged probability exceeded 0.25, on March 21, 2022, was the beginning of the cherry blossom season. The full blossom, or peak, was subsequently confirmed between the 28th and 31st of March 2022. Finally, the blossoms were observed to have dropped with the growth of the leaves on April 7, 2022, as shown by the moving averaged probability falling to 0.25.

**Figure 5.**
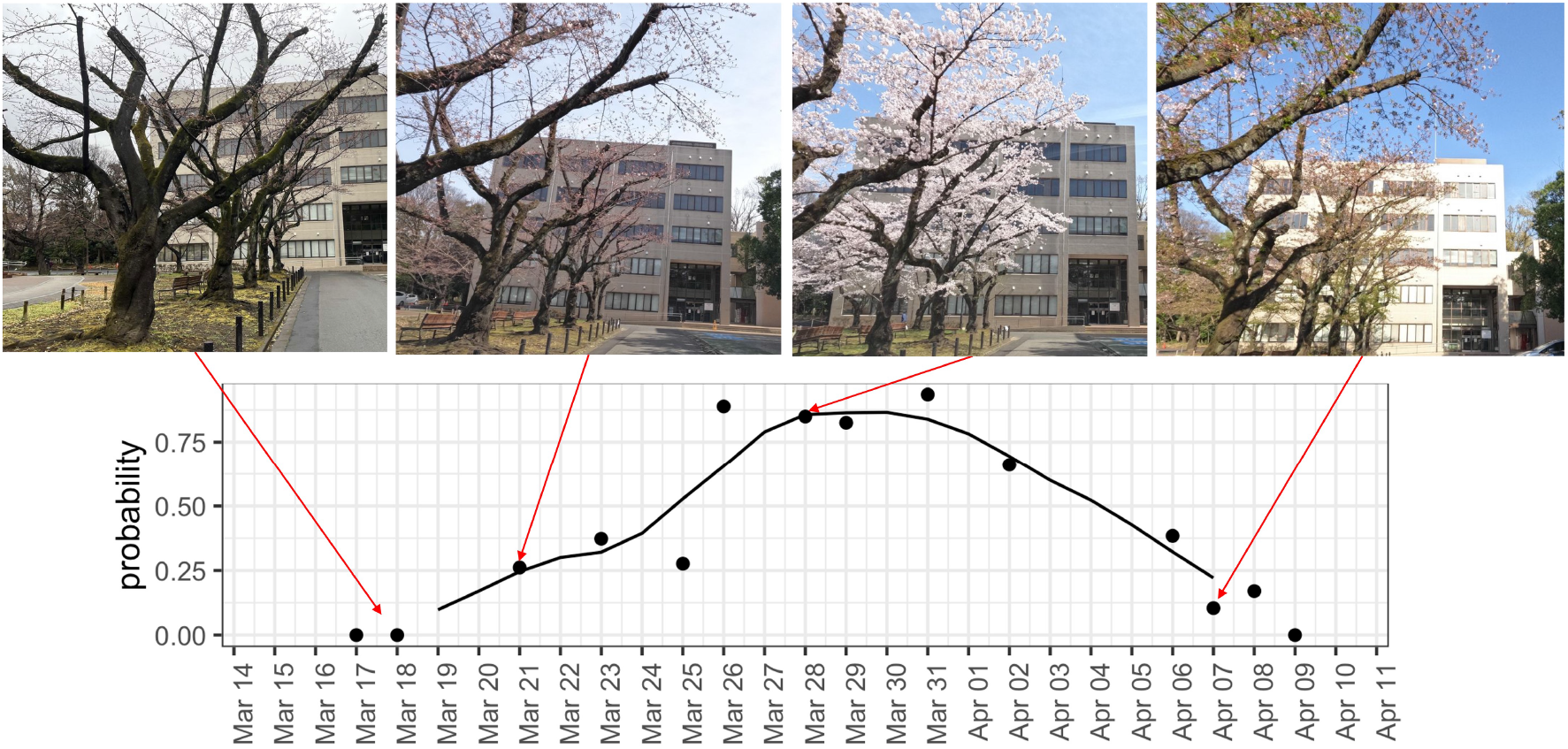
Time series changes in cherry blossoms with four sampled street-level images and calculated detection probability. The dots represent the spatially averaged detection probability of cherry blossoms at a 10 m × 10 m spatial grid, and the line represents the moving averaged probabilities over time.

### 3.2. Accuracy assessment

Two different types of assessment were used to validate the efficacy of our methodology. Firstly, each street-level photograph was run through YOLOv4 model to rigorously evaluate the model’s output and ascertain its accuracy in detecting cherry blossoms. We meticulously selected alternate dates and randomly sampled photographs from each date in an effort to remove any bias resulting from heterogeneous selection across time. A total of 568 photographs were chosen and analyzed to visually determine whether cherry blossoms were accurately detected or not.

Figure 6 shows the histogram of the detection probability for the *true* and *false* samples, which represent cherry blossoms that are present and absent in photographs, respectively. A threshold probability value of 0.25 was deemed suitable for determining cherry blossom detection accuracy. Nevertheless, several *true* samples were identified with a low probability score below this threshold, which implies the existence of false positives. The distribution visualization for the *false* samples, on the other hand, demonstrated good discrimination, with almost all (above 95%) of them having been assigned a probability of 0.25.

**Figure 6.**
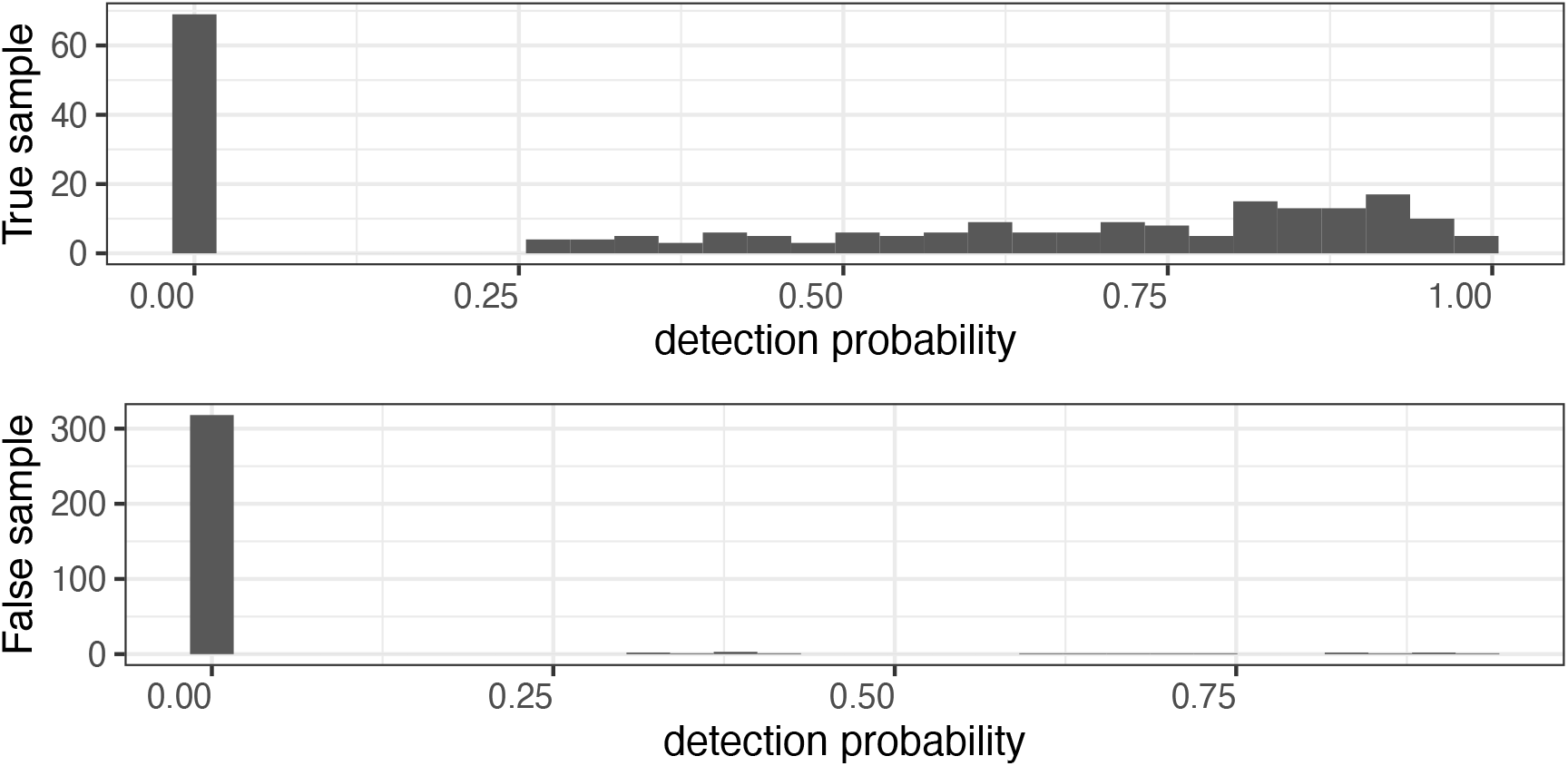
Histograms of detection probabilities for true and false samples.

Therefore, the use of a detection probability threshold of 0.25 was implemented with the objective of achieving a definitive categorization in all images. The results of this assessment showed an 86.7% overall accuracy, coupled with a 70.3% recall and a 90.1% precision, as depicted in Table 1.

**Table 1.** Confusion matrix of cherry blossom detection.

|  |  | Prediction |  |
| --- | --- | --- | --- |
|  |  | True | False |
| Reference | True | 163 | 69 |
|  | False | 18 | 318 |

## 4. Discussion

The proposed methodology aims to map the probability of cherry blossom blooms. The success of this approach is expected to revolutionize the development of semi-automatic phenological observation systems based on social sensing photographic data. We document cherry blossom flowering occurrences, specifically those related to its start, peak, and end stages, using geotagged street-level images. These images were frequently collected and collated on a gridded map, as shown in this case study. Our approach has a distinct advantage over the system proposed by Morishita et al. (2015), as it can be extended to other flowering species with minimal effort. We can easily create spatiotemporal blooming records for different flowering species with minor adjustments in the detection model and by training it with a relatively small body of annotated photos (less than a thousand).

As illustrated in Figure 1, our approach consisted of three essential components: data, model, and phenological analysis. Our methodology prioritizes efficiency and applicability, ensuring its universal applicability across a wide range of locations and flora-related events.

The primary source of concern in the data section is data availability. While the authors were able to take numerous photographs of the study area during their research, the frequency and geographic scope of their observations may be insufficient due to labor and time constraints associated with observation costs.

As a result, implementing a citizen-science methodology may be critical in maintaining an effective observation system. Unlike traditional citizen-science campaigns that collect field observations, our approach does not require the manual input of field data, such as specific degrees of blooming or location and date information. Rather, our approach necessitates the taking of street-level photos with geotags across both time and space. Another advantage of our citizen-science approach is that volunteers do not need to be experts in ecology or remote sensing methods. Instead, it is expected that a wide range of prospective volunteers—including car drivers, hikers, bikers, and walkers— may participate. Many unidentified volunteers will most likely take geotagged street-level photos and upload them to Mapillary to create maps of cherry blossoms blooming elsewhere. Furthermore, replacing the model portion of our research with different object detection models is simple. We acknowledge that various deep-learning object detection models have recently emerged. A comprehensive review of these models is found in Zaidi et al. (2021).

The efficacy of the proposed method is largely dependent on the detection accuracy of the model employed. When identifying cherry blossoms in street-level imagery, our object-detection model achieved an overall accuracy of 86.7%. However, the assessment revealed a relatively low recall rate of 70.3%, which may result in some under-detection of cherry blossoms captured in photographs. However, because of the sequence of photographs collected in Mapillary, this issue may be resolved by incorporating multi-angle observations captured in time-lapse mode. This method compensates for the overlooking issue and allows for better detection performance. Therefore, while measures to improve the model’s accuracy continue, a sequence of photo observations can provide a solution to the problems associated with overlooking cherry blossoms.

Two major parameters must be considered in the phenological analysis section. These parameters include the spatial resolution of the aggregation and the threshold of the spatially aggregated detection probability, both of which are indispensable for distinguishing blooming and conducting phenological analysis. Current research defines the values of the spatial resolution at 10 meters. While conducting experiments, the positional accuracy of volunteered geographic information requires careful analysis.

Although the majority of the literature shows that positional error is less than 10 meters, Menard and Miller (2011) discovered that some studies found higher positional errors ranging from 7 to 13 meters (Menard et al., 2011). Hence, it may be convenient to choose a spatial resolution, such as 15 meters, to increase the dependability of the data. The threshold of the spatially aggregated detection probability was determined using Figure 6. Increasing the threshold value reduces false positives but decreases the number of photos classified as true positives. More case studies would be required to determine precise threshold values. In this study, we used the threshold approach, a commonly employed technique in phenological analysis, to determine the timing of cherry blooming. A threshold value of 0.25 was used to indicate the onset and termination of the blooming period. While the threshold approach has been widely used for this purpose, it is important to note that the optimal threshold value varies significantly across regions. An alternative method for determining the start and end of blooming is to use derivatives, as suggested by previous studies (Beurs and Henebry, 2010; Tsutsumida et al., 2022).

Our results for the start and peak of blooming differed from those of the Japan Meteorological Agency (JMA) for Tokyo. The JMA record for 2022 (https://www.data.jma.go.jp/sakura/data/index.html) indicated that the blooming in Tokyo, located 22 km to the south-east of the study area, began on March 20, and the peak blooming stage occurred on March 27. Similarly, Kumagaya city, located 38 km to the north-west of the study area, recorded the start of blooming on March 24 and the peak on March 30. The first day of blooming is defined by the JMA as the time when at least five cherry buds on a reference tree bloom, whereas the full bloom is reached when over 80% of the buds on the reference tree are blooming (Horikawa et al., 2022; Japan Meteological Agency, n.d.). A promising area for future research is the development of an object detection model to identify flowers at a bud level, which would allow estimation of the start of blooming from street-level imagery. However, to achieve this, a larger dataset of annotated photos is needed to facilitate classification in accordance with the JMA definition.

To demonstrate the efficacy of our approach, additional applications to create comprehensive records of flora across Japan must be investigated. Furthermore, in order to investigate alternative species such as palms, apricots, hydrangeas, and dandelions, our research must continue to evolve. While the Japan Meteorological Agency (JMA) has the capacity to monitor the floral phenology of these species, financial constraints may limit the agency’s ability to conduct continuous observations. As a result, it is imperative to develop a simple and cost-effective method for creating comprehensive records of the phenology of these species. Our current approach is not case-specific and can be generalized for future applications. It can also be customized by changing the object detection model and the spatial resolution of the aggregation grid, as well as the phenological analysis, whenever necessary.

## 5. Conclusions

In this paper, we present a novel method for constructing maps that depict the geographical distribution of cherry blossoms and predict the phenological events associated with their blooming, which includes the precise recording of the date and coordinates of these phenomena using social sensing photo data. We used a popular deep learning algorithm, YOLOv4, to perform efficient cherry blossom detection using geotagged street-level photos obtained from Mapillary, an open-source platform that facilitates collaborative mapping. By combining the applicability of deep learning and social sensing data, we were able to derive invaluable phenological information about cherry blossoms without the need for manual records, thereby overcoming the limitations of traditional citizen-based observation networks and stationary observations. Our method relies entirely on geotagged, street-level photos taken on mobile devices with GPS sensors such as smartphones or dashcams, thereby avoiding any human-based biases in record keeping. Moreover, our methodology is scalable and theoretically unlimited by human resources. While our findings show that our approach is feasible and viable, future research should focus on improving the accuracy of the detection methods and expanding our framework to analyze other species.

## Author contributions

Narumasa Tsutsumida (NT) conceived the ideas and designed the methodology; NT collected the data; Shuya Funada (SF) developed the model; NT and SF analyzed the data; NT led the writing of the manuscript. All authors contributed critically to the drafts and gave final approval for publication.

## Data availability statement

All geotagged street-level photos used in this study can be found in Mapillary. An example is https://www.mapillary.com/app/?pKey=672537624003910.

## Acknowledgments

We appreciate Mapillary users sharing valuable street-level photos. This work is partly supported by National Geographic Asia Labs Fellow and Japan Science, Technology Agency program ACT-X No. 21455106.

